# Rational modulation of plant root development using engineered cytokinin regulators

**DOI:** 10.1101/2024.12.06.627221

**Authors:** Rohan Rattan, Simon Alamos, Matthew Szarzanowicz, Kasey Markel, Patrick M. Shih

**Affiliations:** Joint BioEnergy Institute, 5885 Hollis Street, Emeryville, CA 94608, USA; Environmental Genomics and Systems Biology Division, Lawrence Berkeley National Laboratory, Berkeley, California, USA; Department of Plant and Microbial Biology, University of California, Berkeley, CA 94720, USA; Department of Bioengineering, University of California, Berkeley, California, USA; Innovative Genomics Institute, Berkeley, California, USA

## Abstract

Achieving precise control over quantitative developmental phenotypes is a key objective in plant biology. Recent advances in synthetic biology have enabled tools to reprogram entire developmental pathways; however, the complexity of designing synthetic genetic programs and the inherent interactions between various signaling processes remains a critical challenge. Here, we leverage Type-B response regulators to modulate the expression of genes involved in cytokinin-dependent growth and development processes. We rationally engineered these regulators to modulate their transcriptional activity (*i.e.*, repression or activation) and potency while reducing their sensitivity to cytokinin. By localizing the expression of these engineered transcription factors using tissue-specific promoters, we can predictably tune cytokinin-regulated traits. As a proof of principle, we deployed this synthetic system in *Arabidopsis thaliana* to either decrease or increase the number of lateral roots. The simplicity and modularity of our approach makes it an ideal system for controlling other developmental phenotypes of agronomic interest in plants.

## Introduction

Plant development is controlled by a complex interplay of genetic and environmental cues. Central to this process are hormonal signaling networks, which tightly regulate growth, differentiation, and adaptation in response to changing environmental conditions. Key hormones, such as auxin and cytokinin, serve as central regulators of developmental pathways, orchestrating processes like cell division, elongation, and organ formation^1–3^. As plants continuously adjust to environmental stressors and nutrient availability, these hormone-driven signaling pathways become pivotal in regulating plant phenotypes^4^. In recent years, efforts to manipulate plant development through modulation of these hormonal pathways have gained traction, offering the potential to enhance traits such as yield, stress tolerance, and overall growth vigor^4,5^. Such approaches hold promise in addressing global agricultural challenges, where the ability to fine-tune developmental outcomes can improve plant resilience and productivity under diverse environmental conditions^6^.

Among the various developmental processes influenced by plant hormonal networks, root system architecture stands out due to its fundamental role in nutrient acquisition and water uptake. Root configuration is a critical determinant of a plant’s ability to efficiently explore and exploit the surrounding soil environment, accessing vital resources such as minerals and water. Among the various components of the root system, lateral roots have garnered particular attention, as they serve as key drivers of root branching and contribute directly to nutrient use efficiency and water absorption^7^. The ability to precisely and predictably modulate lateral root formation holds immense potential for optimizing plant performance under diverse environmental conditions seen with the effects of climate change^8,9^. As such, developing systems capable of tuning important phenotypes, like lateral root count, has become an emerging goal in plant biology research.

Lateral root formation is tightly regulated by hormonal pathways, with cytokinin signaling playing a pivotal role in the negative regulation of this process^10^. A key component of the cytokinin signaling pathway involves Type-B response regulators (Type-B ARRs, with “A” denoting *A. thaliana*), a family of transcription factors (TFs) that bind cis-regulatory elements in cytokinin-responsive genes. Upon cytokinin sensing, a cascade of phosphotransfer events is initiated, beginning with histidine kinases (AHKs) that phosphorylate histidine phosphotransfer proteins (HPTs) (Fig. 1A). HPTs activate Type-B ARRs by phosphorylating a conserved aspartate residue in their receiver domain. Type-B ARRs structures consist of Myb-like DNA-binding domains in the C-terminal region and a receiver domain in the N-terminal region. Once activated, Type-B ARRs act as TFs that regulate the expression of cytokinin-responsive genes, including Type-A ARRs, which provide negative feedback regulation to fine-tune the cytokinin response^11–13^. The mechanism of negative feedback still remains undefined, but it is thought to be some combination of competition with Type-B ARRs for phosphotransfer as well as phospho-dependent interactions with target proteins^14^.

**Fig. 1:**
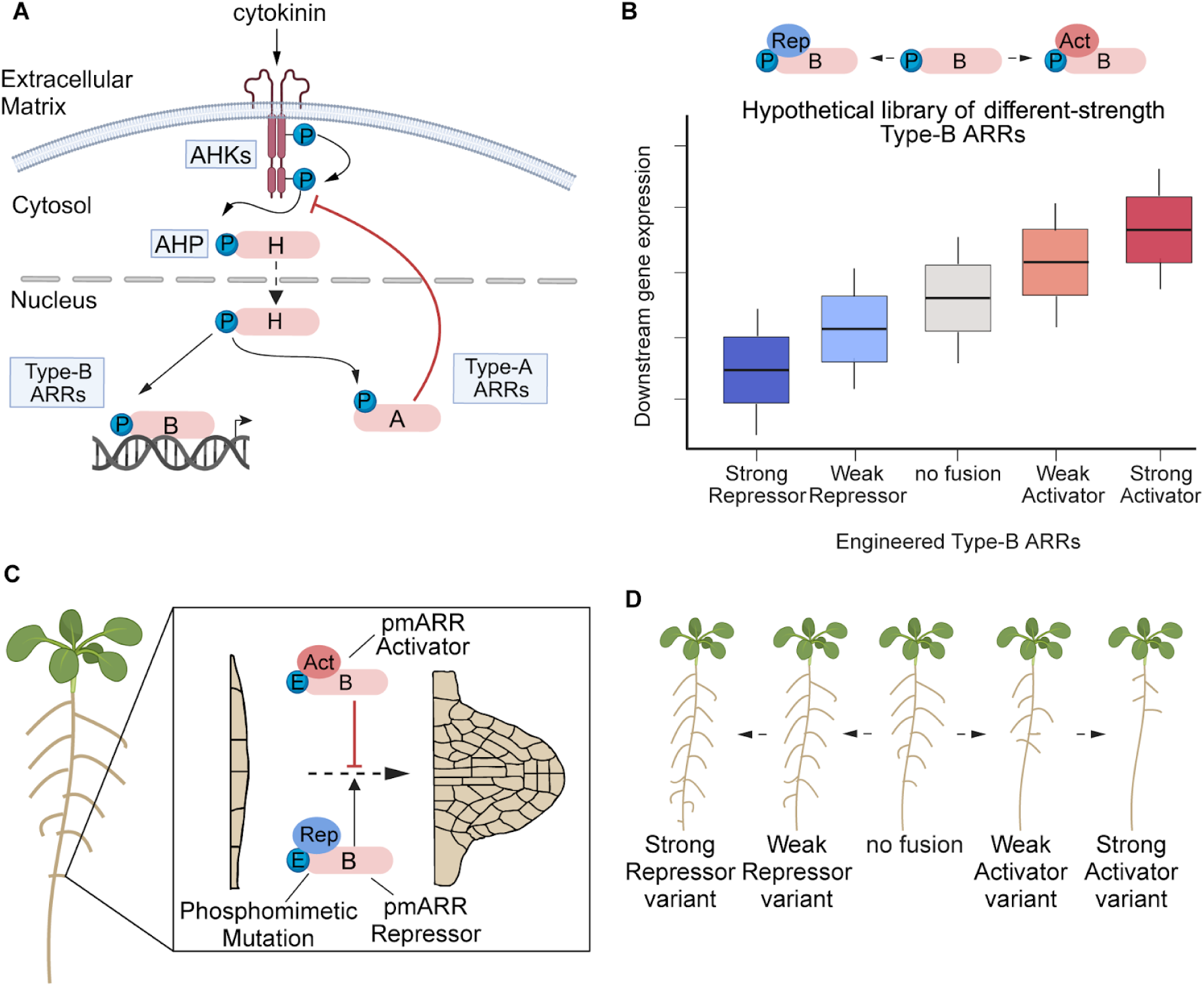
Synthetic biology approach to modulate plant development using cytokinin signaling machinery. **(A)** Diagram of the canonical cytokinin signaling transduction pathway. Cytokinin perception by AHK receptors at the plasma membrane triggers a phosphotransfer cascade, activating Type-B ARRs to modulate transcription. **(B)** Hypothetical schematic of the characterization process for Type-B ARRs of varying regulatory strength. Regulatory strength is quantified by measuring downstream gene expression and achieved by fusing diverse activation and repression domains to a Type-B ARR. **(C)** Figure showing the predicted effects of repressor and activator phosphomimetic Type-B ARRs on lateral root initiation, where an activator phosphomimetic ARR acts as a negative regulator and repressor phosphomimetic ARR as a positive regulator of lateral root development. **(D)** Cartoon illustrating the expected outcomes of deploying an engineered phosphomimetic ARR library to root cells in *A. thaliana*, with the lateral root phenotype scaling in accordance with ARR strength.

Given the central role of Type-B ARRs in mediating cytokinin responses, we selected this family of TFs as amenable candidates for engineering the cytokinin signaling pathway. The approach of fusing regulatory domains to TFs has a strong precedent as a strategy for modulating their activity. It has been demonstrated that fusing the SRDX EAR repression motif to activator TFs can dampen their activity or even convert them to repressors^15^. In addition, previous studies have demonstrated that the activity of activator TFs can be significantly amplified by fusing them with strong activation domains like VP16^16,17^. Furthermore, it has also been shown that tuning the activity of Type-B ARRs is feasible, as demonstrated by the ARR1-SRDX variant, which can effectively suppress cytokinin signaling when overexpressed in *A. thaliana*^18^. Plants overexpressing ARR1-SRDX had a higher density of lateral roots than WT, indicating that the normal role of ARR1 as a transactivator can be reversed to a repressor. However, because of the strength of SRDX and/or the strong and ubiquitous expression of the 35S promoter, this effect was extremely severe and pleiotropic. Additionally, since Type-B ARRs are generally considered to be activators, we hypothesized that it may be possible to enhance cytokinin signaling by fusing Type-B ARRs to activation domains of different strengths and thereby achieve a larger dynamic range of tunability. With this foundation, we aimed to employ this strategy by constructing a library of engineered Type-B ARRs, incorporating both activation and repression domains at the TF’s C-terminus to generate repressor and activator variants with a range of regulatory strengths capable of tuning cytokinin responses (Fig. 1B). ARR12, a member of the Type-B family, was selected as the Type-B of choice due to its well-established role in lateral root formation as well as its function in shoot regeneration, making it an ideal candidate for modulating key developmental processes through the cytokinin pathway^19,20^. ARR12 is typically classified as a weak transcriptional activator, and it contains an acidic activation domain that is sufficient for transactivation in a yeast-based reporter assay^21^. ARR12 and ARR1 act as negative regulators of lateral root formation, functioning redundantly to mediate cytokinin’s inhibitory effect on this developmental process^22,23^.

To overcome the challenges of endogenous regulation, an ideal engineered system requires both orthogonality and dominance over native regulation. Orthogonality is essential to ensure the engineered variants remain active independently of internal cytokinin fluctuations, preventing interference from endogenous signaling. We sought to address this goal by incorporating phosphomimetic variants such as Type-B ARRs with D→E mutations on a conserved aspartate residue within the receiver domain. Previous studies have established a similar strategy, demonstrating that the D94E ARR1 variant induces cytokinin insensitivity by mimicking cytokinin-dependent phosphorylation, effectively bypassing endogenous regulation and providing a blueprint for engineering constitutively active Type-B ARRs^24^. In regards to dominance, many studies have demonstrated that activator TFs fused with the SRDX repression motif exhibit dominant-negative properties^15^. Dominance would ensure the desired phenotypic outcomes without the need for a specific genetic background.

To achieve predictable control over a discrete cytokinin-dependent developmental pathway while minimizing pleiotropic effects, we reasoned that tissue-specific expression of our engineered Type-B ARRs would be critical. For instance, targeting the expression of engineered ARRs to lateral root founder cells could allow direct modulation of lateral root density without disrupting normal functions in other tissues (Fig. 1C). Given the role of cytokinin as a negative regulator of lateral root formation, we anticipate that strong activators expressed in lateral root precursor cells will lead to a diminished lateral phenotype, while strong repressors may increase the number of lateral roots (Fig. 1D).

In this study, we sought to implement this strategy as a proof of principle for the predictable tuning of plant development. We demonstrate that the Type-B ARR ARR12 can be rationally engineered to alter its regulatory function and we test to what extent these engineered variants are insensitive to cytokinin signaling.

## Results

### Design of engineered ARR12 variants with tunable regulatory activity

To generate a comprehensive library of ARR12 variants with varying regulatory logic (*i.e.*, repressors or activators) and strength, we fused 5 repression and 2 activation domains to the C-terminus of the *A. thaliana* ARR12 gene (Fig. 2A). The repression domains included a concatemer of the well-known SRDX synthetic EAR motif, which in turn is based on a peptide from the C-terminus of Arabidopsis SUPERMAN (Fig. 2A). The other 4 repressor domains correspond to previously characterized mutant versions of SRDX (known as KMTR motifs)^25^. The KMTRs were chosen to span a large range of repression strengths based on previously published Gal4 recruitment reporter assays^25^ (Fig. 2A). The activation domains tested included PR, a fusion of the *H. sapiens* P65 and RTA activation domains (originally found as part of the VPR activator, but lacking the VP16 part^26^) as well as the plant-derived EDLL motif, originally from the Arabidopsis ER89 transcription factor^27^. Finally, this 8-member collection also included ARR12 by itself (without any C-terminal fusion, hereafter referred to as -Fusion), which is expected to behave as an activator. To achieve TFs with constitutive activity, we engineered phosphomimetic variants of ARR12, designated as pmARR12, by introducing a D74E substitution in the conserved cytokinin-responsive phosphorylation site^24^.

**Fig. 2.**
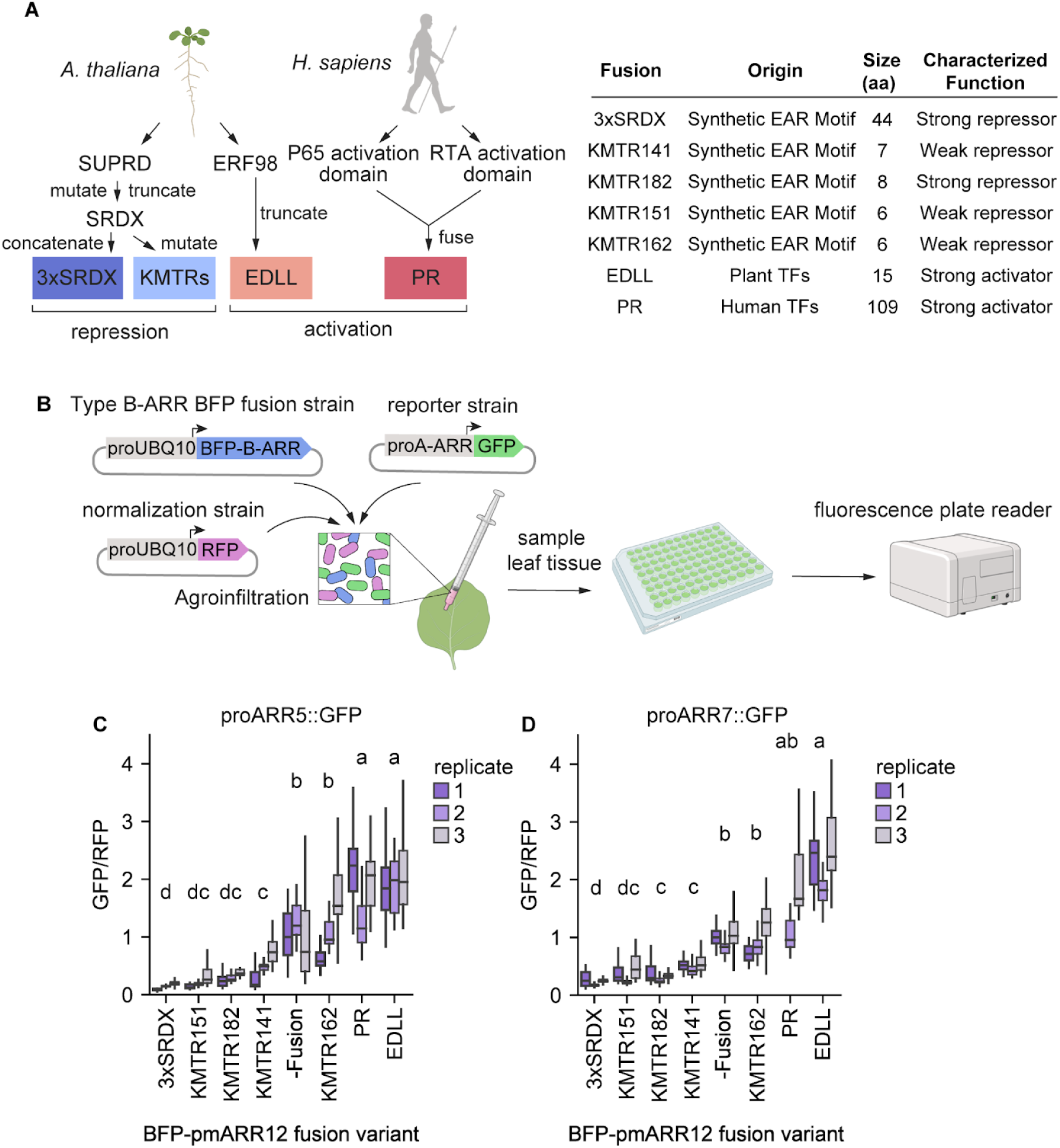
A library of engineered repressor and activator type-B ARRs of varying strengths. **(A)** Left: Origin of the activation and repression domains fused to ARR12 (see text for details). Right: table detailing the origin, total amino acid size, and the previously established function of the various transcriptional domains. **(B)** Schematic depicting the *Agrobacterium*-mediated transgene assay used to characterize engineered Type-B ARRs. The assay involves the coinfiltration into *N. benthamiana* of three *Agrobacterium* strains, each carrying a different binary vector. Effector strains carry engineered variants of Type-B ARRs fused to BFP and driven by the Arabidopsis UBQ10 promoter (proUBQ10). Reporter strains carry GFP under the control of a Type-A ARR promoter regulated by Type-B ARRs. Finally, a strain carrying RFP under proUBQ10 is included for internal signal normalization. These strains are mixed in equal ratios and infiltrated into *N. benthamiana*. Leaf tissue is collected in a microplate and the fluorescence of each construct is measured in a plate reader. **(C)** Box plots showing the normalized GFP output driven by the ARR5 promoter in *N. benthamiana* leaf discs for each BFP-pmARR12 fusion variant in 3 different replicates (n = 36 leaf disks per variant per replicate). **(D)** Same as (C) for the ARR7 promoter. Box plots in (C) and (D) show the median ± IQR, whiskers show the minima and maxima excluding outliers, if present. A Kruskal-Wallis test was run on each dataset to determine if there were significant differences between variants, resulting in P = 2.8160e-89 for ARR5 and P = 7.9566e-108 for ARR7. The letters on top of each variant represent statistically significant groups (P < 0.05) based on Dunn’s test.

To rapidly characterize these variants *in planta*, we utilized the *Agrobacterium*-mediated transient gene expression system in *Nicotiana benthamiana,* which has previously been employed to study the transactivation of Type-A promoters by Type-B ARRs (Fig. 2B)^28^. Specifically, we utilized a three-strain coinfiltration-based system. Effector strains carried a plasmid coding for pmARR12 driven by the strong *A. thaliana* UBQ10 promoter (Fig. 2B). These pmARR12 variants were fused to a blue fluorescent protein (BFP) at the N-terminus to measure the pmARR12 expression level and study the role of TF dosage in our assay. The second coinfiltrated strain carried a reporter gene composed of either the ARR5 or ARR7^29^ promoters from *A. thaliana*, driving the expression of GFP and providing a quantitative readout of transcriptional activity in response to the different pmARR12 constructs (Fig. 2B). These ARR5 and ARR7 promoters were defined as 2 kb upstream of the start codon, including the 5’ UTR, following a previous approach that was shown to capture sufficient cis-regulatory elements for detectable promoter activity^30^. Treatment of Arabidopsis roots with exogenous cytokinin induces changes in transcript abundance for hundreds of genes, not all of which respond with the same speed, in the same direction, or to the same degree.^29^ We reasoned that including two cytokinin-responsive promoters would provide a more comprehensive picture of this regulatory diversity. The ARR5 and ARR7 promoters contain the canonical AGATHY cis-elements, a sequence motif directly bound by Type-B ARRs like ARR12, enabling the direct regulation and activation of these reporters by ARR12^31,32^. Finally, to account for the variation in GFP fluorescence that is not due to upstream ARR12 expression, we coinfiltrated a control strain carrying RFP under the UBQ10 promoter. Dividing the reporter signal by that of a constitutive transgene is a standard practice in the field that is meant to account for experimental error affecting expression of both the reporter and normalization transgenes in a similar fashion.

### Characterization of engineered ARR12 variants in *N. benthamiana*

We infiltrated *N. benthamiana* with 3-strain mixes containing each of the eight engineered pmARR12 variants, the ARR5/7 GFP reporter and the RFP normalization strain. As a baseline control for the reporter activity, we used the empty vector backbone (hereafter EV) instead of ARR12 (hereafter referred to as the no ARR12 infiltration). To better understand what aspects of the data were reproducible and to what extent, we repeated these experiments in three different weeks using different batches of plants. Measurement of BFP, GFP, and RFP fluorescence allowed the simultaneous determination of pmARR12 expression level, reporter expression, and normalization signal strength, respectively (Fig. S1A).

Examining the level of expression of the RFP normalization transgene revealed that the no ARR12 infiltration infiltration mix carrying EV instead of ARR12 had a much higher fluorescence than the rest, suggesting that expression of ARR12 has a negative, nonspecific effect on transgene expression (Fig. S1A). To explore this possibility, we coinfiltrated the -Fusion BFP-pmARR12 variant with various constitutively expressed RFP and GFP normalization transgenes driven by three different promoters, using EV as a control (Fig. S1B). Across all combinations, -Fusion reduced GFP and RFP fluorescence intensity by approximately 10-fold compared to no ARR12, showing that the expression of effector pmARR12 constructs represses transgene expression in *N. benthamiana*, independent of transgene sequence (Fig. S1B). Nonetheless, without considering the no ARR12 mix, the levels of RFP fluorescence were relatively consistent across pmARR12 variants (Fig. S1A). Thus, while RFP normalization cannot be used to compare a given pmARR12 variant with the no ARR12 control, this approach can still be used to account for experimental variation among pmARR12 variants.

We next wondered if the BFP signal could be used to account for part of the observed experimental variation. Transcription can be sensitive to the concentration of transcription factors and we found some consistent differences in pmARR12 expression across variants (as measured by BFP fluorescence). It is thus conceivable that part of the differences in GFP reporter expression are due to differences in pmARR12 abundance. To test if this was the case in our experimental setup, we titrated the -Fusion variant by varying the OD (Agrobacterium optical density at 600 nm) of this strain while keeping the OD of the ARR5 reporter strain constant, and the total OD of bacteria constant using EV (Fig. S1C). The concentration-dependent effect of the -Fusion variant on the ARR5 promoter was extremely weak (Pearson r = 0.12 for ODs 0.05 and higher) and plateaued at a relatively low OD of about 0.05, much lower than the OD used in our ARR12 library characterization experiments (OD 0.15) (Fig. S1C). Furthermore, the pmARR12 variant expressed at the lowest level had a BFP fluorescence value comparable to the saturation BFP value in our titration experiment (Fig. S1C). This suggests that our variant comparison experiments operated in a saturation regime where variation in BFP-pmARR12 abundance has little effect on reporter expression. In agreement with this, we did not find a consistent correlation between BFP and GFP fluorescence across leaf punches (Fig. S1D). In contrast, there was a good and reproducible correlation between the normalization RFP strain and the reporter GFP expression, confirming that RFP normalization can be used to compare between variants (Fig. S1D). Given these results, we used normalized GFP/RFP for comparisons between variants and raw GFP to compare variants to the no ARR12 reporter alone infiltration.

Comparing the levels of normalized GFP fluorescence driven by the ARR5 reporter mixed with different pmARR12 fusions revealed significant differences in activity across engineered variants. As expected, the -Fusion variant, as well as PR and EDLL acted as activators of the ARR5 promoter compared to no ARR12 (Fig. S1A, Fig. 2C). PR and EDLL were stronger activators of the ARR5 promoter than -Fusion, validating our approach to engineer enhanced Type-B ARRs (Fig. 2C). Unexpectedly, the EAR motif variant KMTR162, which acts as a repressor when fused to the Gal4 DNA binding domain, behaved as an activator when fused to pmARR12, showing that there are limits to the modularity of repression motifs. This observation highlights the need to empirically test regulatory domains (Fig. S1A). The rest of the EAR motifs acted as repressors of various strengths (Fig. S1A). KMTR141, being a weak repressor, turned ARR12 into a neutral TF with regards to ARR5 expression (Fig. S1A). These trends were largely the same for the ARR7 promoter, with some exceptions. For this promoter, the -Fusion variant did not increase GFP fluorescence compared to the no ARR12 infiltration (Fig. S1A). Fusing pmARR12 to regulatory domains was necessary for the TF to behave as an activator of the ARR7 promoter (Fig. S1A). All EAR motif fusions acted as repressors of ARR7, but, unlike for ARR5, the KMTR141 variant became a true repressor rather than a neutral TF. The differences between ARR7 and ARR5 are not entirely surprising given the diverse responses observed for these and other cytokinin-inducible genes upon hormone treatment or genetic perturbations^18^. Testing multiple promoters enables a more comprehensive assessment of how engineered TF variants influence target gene expression.

In sum, we were able to characterize a library of pmARR12 constructs with statistically significant differences in functional strength (Fig. 2C,D, Fig. S1A). This effort established a collection of cytokinin regulator TFs with varying repressive and activating potentials capable of altering the expression profile of native *A. thaliana* promoters.

### Modulation of lateral root development in A. thaliana using engineered Type-B ARRs

Next, we sought to test if quantitative differences in regulatory activity across engineered ARR12 variants, as measured by agroinfiltration, can predict differences in a quantitative phenotype controlled by cytokinin. To this end, we tested whether transgenes carrying 3 of our engineered variants could predictably modulate lateral root density in stably transformed *A. thaliana*. To mitigate the risk of pleiotropic effects that could arise from ectopic expression of engineered TFs, we employed tissue-specific promoters. The cell type-specific promoter *proLBD16* was initially chosen for its established roles in lateral root development. The LBD16 gene is expressed during the early stages of lateral root primordium initiation^33^. By driving the expression of our engineered pmARR12 variants under this promoter, we aimed to confine the effects to lateral root formation, minimizing off-target effects and ensuring a focused developmental readout. We selected representative constructs from our library, including a strong activator (PR), a weak activator (-Fusion), and a strong repressor (3xSRDX).

Next, we generated at least 10 stable transgenic lines in a wildtype (WT) *A. thaliana* (Col-0) background carrying the selected BFP-pmARR12 variants under the *proLBD16* promoter. We then used T2 seedlings to determine lateral root count and primary root length, with lateral root density defined as the ratio between these two metrics (see Materials and Methods). As cytokinin signaling negatively regulates lateral root formation and growth, we expected the strong repressor to increase lateral root density compared to the weak and strong activator variants. Following the same rationale, we expected the strong activator to have fewer lateral roots than the weak activator. Initial results with the proLBD16 transgenic lines were inconclusive; the engineered regulators did not significantly impact the lateral root phenotype, and the lateral root densities among the three variants were not significantly different (Fig. 3A). This suggested that proLBD16-driven expression may not provide sufficient sensitivity to enable detection of subtle phenotypic changes in lateral root formation. Additionally, it is plausible that the proLBD16 promoter lacks the necessary spatiotemporal expression pattern to influence this phenotype effectively. If LBD16 is primarily expressed in the primordium, its activation may occur too late to impact lateral root initiation.

**Fig. 3:**
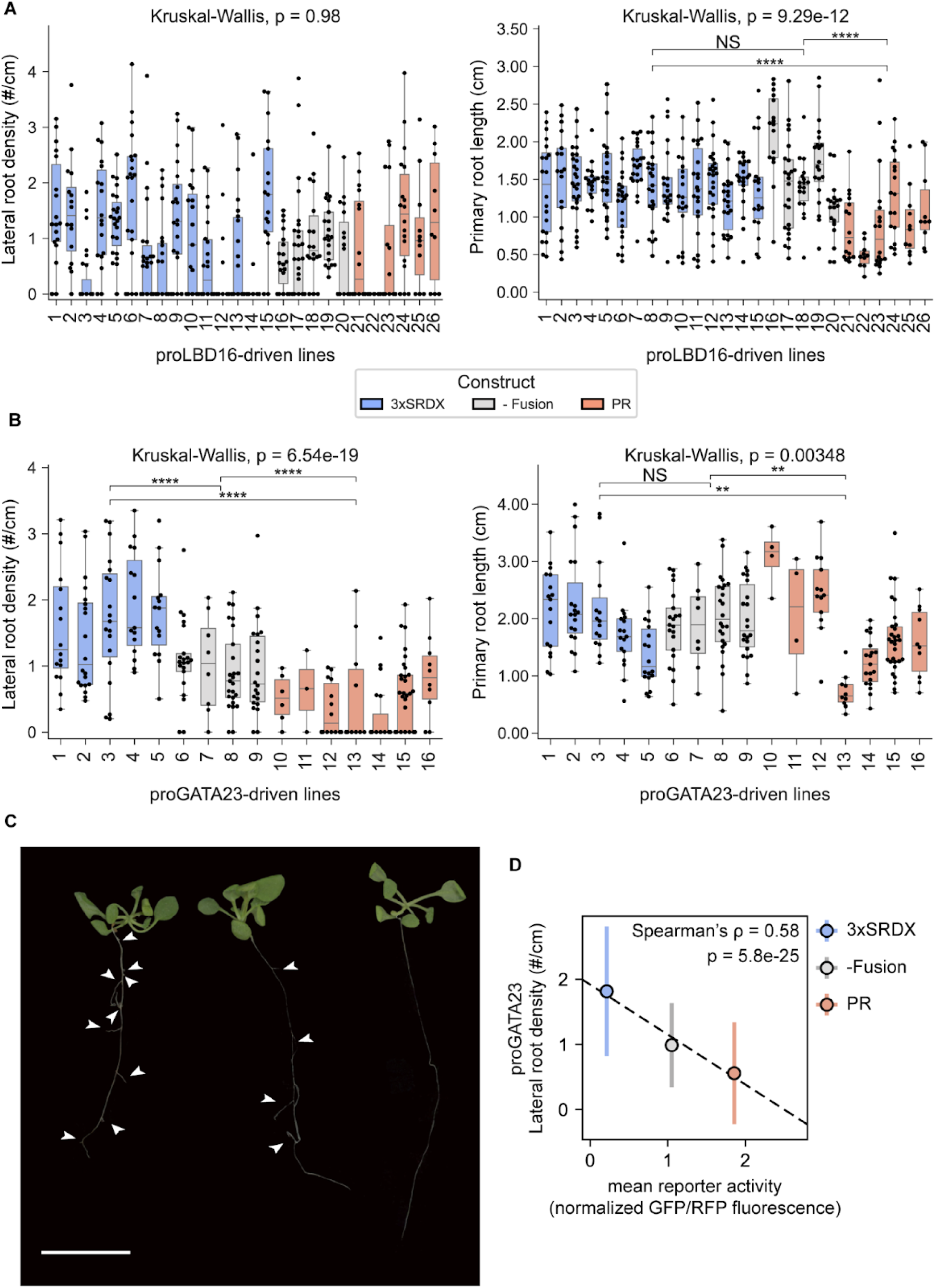
Bidirectional and predictable modulation of lateral root density by cell-type specific expression of pmARR12 variants. **(A)** Left: Box plot of lateral root density (lateral root count / primary root length) of T2 transgenic *A. thaliana* individuals with either a repressor, activator, or no fusion pmARR12 variant driven by the root-specific promoters proLBD16. Right: Box plots of the primary root length of the same set of constructs and lines. **(B)** same as (A) for the proGATA23 promoter. A Kruskal-Wallis test comparing each distinct plant genotype to one another was used to determine the effect of the transgene in each phenotype. NS (not significant, P > 0.05), *P < 0.05, **P < 0.01, ***P < 0.001, and ****P < 0.0001. **(C)** Image showing*A. thaliana* seedlings engineered to modify root branch density, using proGATA23-driven variants: a proGATA23::BFP:pmARR12:PR activator line (Line 12, right), a proGATA23::BFP:pmARR12 line (Line 7, middle), and a proGATA23::BFP:pmARR12:3xSRDX repressor line (Line 4, left). Arrowheads indicate visible emerged lateral roots. Scale bar = 1 cm. **(D)** Mean ± SD lateral root density (y axis) as a function of TF variant strength (x axis, combining all data from both promoters in Fig. 2C and D) for the variants tested in Arabidopsis. The dashed line shows a linear fit to the data.

Motivated by these findings, we hypothesized that targeting an earlier stage in root development might yield more pronounced effects. Therefore, we subsequently employed the proGATA23 promoter, which regulates the expression of GATA23, a key TF known to participate in the specification of lateral root founder cells^34^. In contrast to the proLBD16 lines, we observed a significantly higher number of lateral roots in the 3xSRDX line compared to the activator lines. Furthermore, the strong PR activator line had significantly lower lateral root density than the weak activator -Fusion line (Fig. 3B,3C). The median primary root length did not change between the -Fusion lines and the repressor lines, but median primary root length across PR lines was 30% shorter than the repressor lines and 22% shorter than the control lines (Fig. 3B,3C). Importantly, these differences in primary root lengths do not explain differences in lateral root density since a shorter primary root would result in an overestimation of the lateral root density. Thus, the density measurement reflects the true modulation of lateral root formation rather than being an artifact of altered primary root length. Notably, the transcriptional activity of each variant from the tobacco reporter assay was found to be predictive of their respective phenotypic effects (Spearman’s ⍴ = 0.58), validating our pipeline (Fig. 3D).

In prior studies, 35S::ARR1:SRDX transgenic lines reported severely stunted growth within the shoot architecture^18^. In contrast, our engineered lines maintained normal shoot morphology, maintaining structural integrity across the shoot regions (Fig. 3C). Notably, the effects of our engineered TFs were largely confined to the intended target tissues, demonstrating effective spatial specificity.

We next took advantage of the BFP fluorophore fused to pmARR12 to determine the protein expression pattern of the engineered TF in the proLBD16 and *proGATA23* transgenic lines. Nuclear BFP was undetectable in two proLBD16::BFP:pmARR12:3xSRDX lines (Fig. 4A). In proGATA23::BFP:pmARR12:3xSRDX lines, we observed clear nuclear BFP fluorescence in lateral root primordia and emerged lateral roots of the 3xSRDX fusion line, but this signal weakened as lateral roots matured (Fig. 4B, S2). This protein expression pattern was somewhat surprising since transcription of the *proGATA23* promoter is known to be restricted to just a few lateral root initial cells. This discrepancy could be the result of post-transcriptional dynamics regulating the accumulation of BFP-pmARR12. Indeed, GUS reporter assays have shown that in WT plants the ARR12 protein accumulates in young emerging lateral roots, even though its promoter is not active in these cells^19,35^. These dynamics may partially explain why this transgene was effective at regulating lateral root formation and growth. Although proGATA23 is active within a narrow developmental window, its early expression in lateral root founder cells is sufficient to initiate changes in lateral root density.

**Figure 4.**
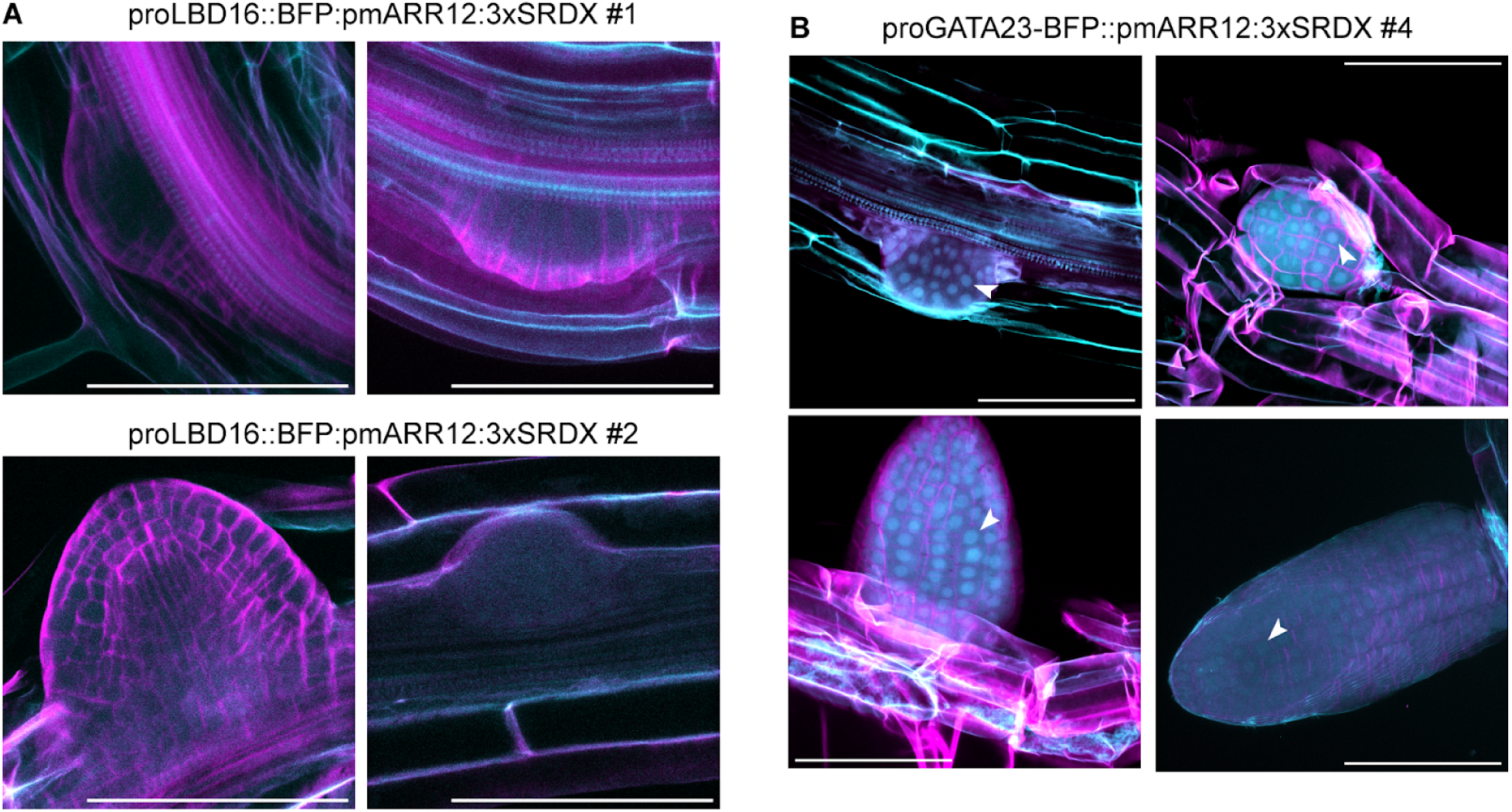
Localization of engineered ARR12 variants in emerging lateral roots. Nuclear BFP was detected in proGATA23 roots but not in proLBD16. **(A)** Maximum intensity projections of confocal fluorescence image stacks of lateral roots at different developmental stages from transgenic lines 1 and 2 of proLBD16::BFP:pmARR12:3xSRDX, stained with the cell wall dye Direct Red 23. The cyan channel corresponds to BFP fluorescence and cell wall autofluorescence. Magenta corresponds to stained cell walls. **(B)** Identical to (A), but for proGATA23::BFP:pmARR12:3xSRDX Line 4. Arrowheads indicate lateral root nuclei containing nuclear-localized BFP. Images were obtained from separate individual T2 seedlings derived from the same T1 line. Scale bar = 100 µm.

### Engineered TF sensitivity to external perturbations

Our engineered ARR12 constructs were sufficient to modulate lateral root density when expressed in a WT background, demonstrating a high degree of dominance. This data suggests that these proteins are relatively insensitive to endogenous changes in cytokinin concentration, as expected for phosphomimetic Type-B ARRs. To challenge the performance of our engineered system, we sought to test whether these alterations to root morphology persist under severe exogenous perturbations. It is well established that cytokinin addition inhibits lateral formation and main root elongation in WT seedlings^36,37^. Hence, to test the robustness of engineered variants, we asked whether the proGATA23::BFP:pmARR12:3xSRDX construct was sufficient to increase the number of lateral roots in seedlings grown in the presence of exogenous cytokinin, compared to WT Col-0 plants. Several lines of evidence indicate that transgenes that overexpress Type-B ARRs are capable of overriding the phenotypic effects of exogenous cytokinin. Treatment of detached leaves of WT plants with cytokinin delays senescence, but this effect was largely lost in plants overexpressing ARR1-SRDX^18^. Similarly, ARR1-SRDX overexpressor lines were much less sensitive to root growth inhibition by exogenous cytokinin than WT^18^. Finally, and more directly related to our goals, it was shown that ARR1-SRDX overexpression could partially rescue the loss of lateral roots under concentrations as high as 0.1 uM of the cytokinin analog BA. To drive the Type-B ARR transgene, all these experiments used the 35S promoter, which is expressed at very high levels throughout the plant. It was therefore not clear whether our construct specifically targeting lateral roots would show similar insensitivity to exogenous cytokinin. The phosphomimetic mutation may enhance the overexpression effect, as seedlings overexpressing a phosphomimetic version of ARR1 mimicked the phenotype of WT treated with cytokinin, while overexpression of WT ARR1 did not ^24^. Although the precise molecular mechanism remains to be clarified, it is plausible that the phosphomimetic mutation locks Type-B ARRs proteins in a constitutively active state, thereby uncoupling them from the typical cytokinin signaling cascade. This sustained activation likely disrupts normal cytokinin-mediated regulation^23^.

To test the cytokinin sensitivity of different genotypes, we grew seedlings of WT and one of the transgenic lines carrying the strong repressor construct proGATA23::BFP:pmARR12:3xSRDX in the presence of the cytokinin analog BA (see Materials and Methods). To better characterize the cytokinin sensitivity of these two genotypes, we used five BA concentrations: 5 μM, 1 μM, 0.5 μM, 10 nM, and 5 nM (Fig. 5A,B). As a baseline, we also grew these two lines without added BA. At low cytokinin concentrations (5nM, and 10nM), both genotypes were able to initiate lateral roots (Fig. 5A). Notably, under these conditions, proGATA::BFP:pmARR12:3xSRDX seedlings consistently exhibited significantly higher lateral root density than WT, indicating that the engineered repressor promotes lateral root formation even in the presence of low levels of exogenously delivered cytokinin (Fig. 5A). However, the lateral root density of the transgenic line decreased with respect to the no BA control to a similar extent than WT. Seedlings grown under BA concentrations of 0.5 μM or higher were severely stunted, showed minimal root growth and no lateral roots, regardless of the genotype (Fig. 5A,B).

**Figure 5:**
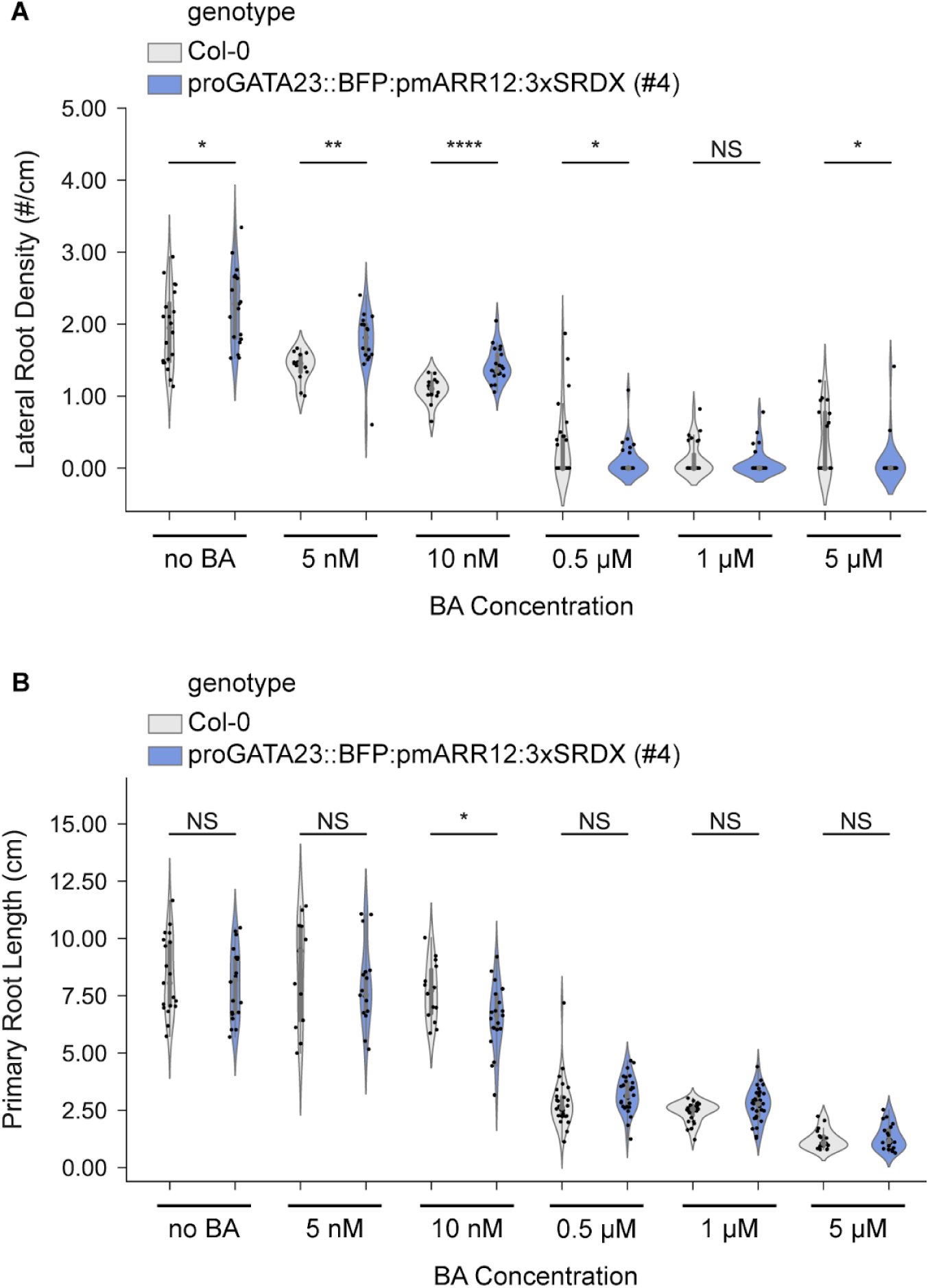
Engineered variant sensitivity to exogenous cytokinin. The repressor line has a higher LR density than WT, but its sensitivity to cytokinin is similar. **(A)** Violin plots showing the lateral root density (lateral root count / primary root length) of *A. thaliana* Col-0 WT and proGATA23::BFP:pmARR12:3xSRDX seedlings grown in varying concentrations of the cytokinin analog BA. T2 individuals from line #4 were used for the transgenic line. **(B)** Violin plots showing the primary root length of the plants shown in a. Bars on top show the result of a two-sided t-test performed between WT and the transgenic line for each BA concentration. NS (not significant, P > 0.05), *P < 0.05, **P < 0.01, ***P < 0.001, ****P < 0.0001.

To confirm that the observed differences in lateral root density are primarily due to changes in lateral root count, we also measured primary root lengths across all tested BA concentrations (Fig. 5B). With the exception of the 10 nM treatment, where a significant reduction was observed in the transgenic line, there were no significant differences in primary root lengths between WT and proGATA::BFP:pmARR12:3xSRDX. This reinforces the conclusion that differences in lateral root density primarily reflect true changes in lateral root initiation, rather than being a secondary effect of altered primary root growth. Note that the primary root lengths reported in this experiment are considerably longer than those of Fig. 3. We attribute this discrepancy to different growth protocols between these two experiments, in particular the use of hygromycin in the growth media in Fig. 3 (Materials and Methods).

These results suggest that, while the proGATA23::pmARR12:3xSRDX construct can enhance lateral root production even under moderate cytokinin levels, its capacity to buffer against cytokinin-induced inhibition is limited. Indeed, it appears that the inhibitory effect of BA is comparable between WT and the transgenic line, but the latter starts at a basal state of higher lateral root density.

One explanation for this observation is that robust root branching requires sustained inhibition of cytokinin signaling in stages of lateral root formation during which the proGATA23-driven engineered ARR12 is not present at high enough levels. The engineered ARR12 most likely accumulates during a limited developmental window (Fig. 4) where it is sufficient to promote lateral root growth (Fig. 3), however it might not be present at high enough levels during earlier stages when pericycle cells commit to the lateral root fate. This contrasts with transgenes driven by a strong and constitutive promoter such as 35S, which are able to exert their function in all cells throughout development.

## Discussion

In this study, we engineered a library of phosphomimetic ARR12 variants with tunable regulatory logic and strength to fine-tune a quantitative developmental phenotype in plants. By fusing activation and repression domains to the C-terminus of the ARR12 TF and expressing them with cell type-specific root promoters, we developed a system capable of finely controlling cytokinin signaling pathways *in planta*. Characterization of these variants in N. benthamiana is critical since the activity of a given regulatory domain when fused to one DNA binding domain (e.g. Gal4) does not necessarily predict its effect when fused to ARR12 as exemplified by KMTR162. Using the modulation of lateral root development in *A. thaliana*, we tested to what extent our system meets two key engineering goals: dominance and insensitivity to cytokinin. We found the transgenes to have a high degree of dominance, as the engineered variants exert their effects independently of the native signaling components, regardless of the genetic background. On the other hand, although the engineered variants carry the phosphomimetic mutation that renders Type-B ARRs insensitive to cytokinin, exogenous BA did reduce lateral root density in one of these transgenic lines. This suggests that spatiotemporal expression patterns and/or expression levels need to be optimized to achieve insensitivity to exogenous cytokinin.

Recent advances have established a robust foundation for a new generation of plant developmental engineering. For instance, Brophy et al. (2022) achieved controlled suppression of root branching through cis-regulatory designs that incorporated buffer logic of varying strengths^38^. In a similar theme, Khakhar et al. (2018) designed hormone-activated Cas9-based repressors (HACRs) capable of regulating hormone-responsive genes, ultimately allowing for the precise control of key developmental traits like shoot branching^39^. Furthermore, Decaestecker et al. (2019) developed CRISPR-TSKO, a modular system for tissue-specific gene knockout that was capable of disrupting lateral root initiation, root cap development, and stomatal lineage differentiation.^40^ Our simple and modular approach complements these plant engineering efforts.

The conservation of cytokinin signaling in plants^41,42^, combined with the modular nature of our engineered Type-B ARR system supports its broad applicability across plant species and developmental processes. Our framework could easily be adapted to specifically target other cytokinin-mediated developmental processes by substituting the tissue-specific promoters driving effector expression. This adaptability provides a valuable tool for fine-tuning other plant developmental processes beyond lateral root regulation. For example, cytokinin signaling regulates shoot meristem size via the direct activation of WUSCHEL (WUS) by Type-B ARRs^43–45^. In turn, the levels and spatial pattern of WUS expression dictate the size and shape of fruits and other reproductive organs. It may thus be possible to fine-tune these phenotypes with engineered Type-B ARRs.

Regarding these potential applications, we note that different promoters can drastically affect the phenotypic effect of our engineered TFs, even when their expression largely overlaps in space and time such as proLBD16 and proGATA23. Further, in some cases, tissue and cell type-specificity may be dictated by cytokinin signaling itself. This underscores the importance of empirically testing several promoters. Translational fusions to the TF of interest such as our BFP-ARR12 fusion will make it possible to better characterize these expression dynamics.

## Materials and Methods

### Plant Material and Growth Conditions

*N. benthamiana* was grown following a previously published lab protocol^30^. Plants were grown in an indoor growth room at 25°C and 60% humidity using a 16/8 hour light/dark cycle with a daytime PPFD of ∼120 μmol/m2s. The soil consisted of Sunshine Mix #4 (Sungro) supplemented with Osmocote 14-14-14 fertilizer (ICL) at 5mL/L and agroinfiltrated 29 days after seed sowing.

### Plasmid Construction & Transformation of *A. thaliana*

Plasmids were constructed using Gibson assembly^46^ or Golden Gate assembly^47^, following established protocols. For every construct containing the ARR12 gene, we used the genomic sequence of ARR12 in which all the introns except for the first one were removed. Retaining the first intron was necessary since constructs carrying the ARR12 cDNA displayed toxicity in E. coli.

For plant transformation, the floral dip technique was followed using an established protocol^48^. Seeds from the transformed plants were screened on plates containing 35 mg/L hygromycin. The plates, sealed with micropore tape, contained a medium composed of 1.5% plant tissue culture agar and half-strength Murashige and Skoog (MS) basal salt mixture with nutrients, adjusted to a pH of 5.7. Seedlings were cultivated for two weeks under axenic conditions in a Percival growth chamber set at 24 °C with constant light. After this period, transformed seedlings could be distinguished from non-transformed ones based on their larger size and successful germination.

### *N. benthamiana* agroinfiltration assay

Binary vectors were introduced into *A. tumefaciens* strain GV3101. Colonies were inoculated in liquid medium with the appropriate antibiotics the evening before the experiment. Cultures were grown to an OD600 of 0.8–1.2 prior to resuspension in infiltration buffer (10 mM MgCl2, 10 mM MES, and 200 μM acetosyringone (added fresh), pH 5.7)^49^. Cultures were incubated for 2 hours at room temperature and then combined in equal ODs (final OD600 = 0.15 per strain, 0.45 total OD of bacteria). Each coinfiltration mix included the given reporter strain (Type-A promoter-GFP), an effector strain (Type-B pmARR12 variant), and a normalization RFP strain. For control, an empty vector strain carrying the pCambia1300 backbone was infiltrated at an OD of 0.45. For a baseline of reporter output, each coinfiltration mix included the given reporter strain, an empty vector strain, and the RFP strain each at an OD of 0.15. Leaves 6 and 7 of 4-week-old *N. benthamiana* plants were subsequently syringe infiltrated with the *A. tumefaciens* mixed suspensions. Following infiltration, *N. benthamiana* plants were kept under the same growth conditions as previously described.

For fluorescence measurements, leaves were harvested three days post-infiltration. For each combination, two leaves per plant and three plants per construct were used. Thirty-six leaf disks were collected per condition using a standard 6-mm hole puncher. The disks were then placed abaxial side up on 300 μL of water in 96-well microtiter plates, and fluorescence was measured for GFP (Ex.λ = 488 nm, Em.λ = 520 nm), RFP (Ex.λ = 532 nm, Em.λ = 580 nm), and BFP (Ex.λ = 381, Em.λ = 445) using a Synergy 4 microplate reader (Bio-tek).

Normalization of GFP fluorescence was performed by dividing the raw fluorescence units in the GFP channel by those in the RFP channel, for each leaf disk.

### Lateral Root Assay

Seedlings were vapor-phase sterilized as previously described^50^. Sterilized T2 seeds were plated on 1.5% agar half-strength Murashige and Skoog (MS) medium containing 35 mg/L hygromycin, adjusted to a pH of 5.7. Plates were sealed with micropore tape and were placed vertically in a Percival growth chamber set at 24 °C with constant light. 15 seedlings per plate were grown under these conditions for 17 days prior to data acquisition to allow adequate initiation and development of lateral roots.

For the cytokinin sensitivity assays, T2 seedlings of one repressor line (pmARR12-3xSRDX #4) were grown on an initial set of MS plates under previously described conditions and then transferred to the final plates. WT seedlings (Col-0) were initially plated on MS plates without antibiotic selection and the transgenic line was initially plated on Hygromycin plates. Both lines were stratified for 2 days and then taken out (day 0) and allowed to germinate in the growth chamber for 4 days. Next, seedlings were transferred to either MS plates supplemented with 5 µM 6-benzylaminopurine (6-BA) or control MS plates (with no antibiotic selection), maintaining the same growth conditions. For each plate used in the cytokinin insensitivity assay, 10 sterilized seedlings were sown. Following an additional 13 days, seedlings were imaged for subsequent analysis.

### Whole plant Imaging & Lateral Root characterization

To visualize and quantify emerged lateral roots, plates and plants were imaged using an Epson Perfection V600 photo scanner to capture high-resolution images. Lateral roots were identified and counted under the conditions that they were clearly visible at the macroscopic level. Measurements of primary root length and lateral root number were conducted using ImageJ software (version 1.53k) to ensure precise analysis and reproducibility.

### Fluorescence Microscopy

17 day old T2 seedlings of proGATA23::BFP:pmARR12:3xSRDX transgenic lines were grown as described in Materials and Methods: Lateral Root Assay. The seedlings were fixed in 4% paraformaldehyde and cleared for 2 days using ClearSee following a previously published protocol^51^. Cell walls were stained in a 0.1% Direct Red 23 ClearSee solution for 30 minutes and washed in ClearSee before imaging. For BFP fluorescence 405 nm excitation and 420-475 nm emission was used, which also captures cell wall autofluorescence. Direct Red 23 was imaged with 560 nm excitation and 580 - 620 nm emission.

## Acknowledgements

We thank J. Dalton for the stellar maintenance and management of our growth room. We are also grateful to Amy Lanclot for providing feedback on the manuscript. Figure cartoon representations were sourced from BioRender.com. This work was part of the JBEI (https://www.jbei.org) supported by the US Department of Energy (DOE), Office of Science, Office of Biological and Environmental Research, through contract DE-AC02-05CH11231 between Lawrence Berkeley National Laboratory and the DOE.

## Author Contributions

R.R., S.A., and P.M.S. conceptualized the research. R.R., S.A., M.S., and K.M. contributed biological research materials. K.M. contributed preliminary unpublished data. R.R. and S.A. performed experiments. R.R. and S.A. wrote the code to analyze the data. R.R. and S.A. analyzed the data. RR, SA, and PMS wrote the manuscript. PMS provided all research funding.

## Data Availability

All source data is available from a public Google Drive folder: https://drive.google.com/drive/folders/1GiW7Q-K35BqXnhbzl-fyZkYZlMw4FW0H?usp=sharing The sequences of all plasmids used in this study can be accessed from a public Google Drive folder: https://drive.google.com/drive/folders/1TqtH0v7dW-ZzS5UUrr16NX9YhES--GZP?usp=sharing The code used to analyze the data and generate the figures is described via Github at https://github.com/shih-lab/Root_Engineering

## Supporting Information

**Fig. S1:**
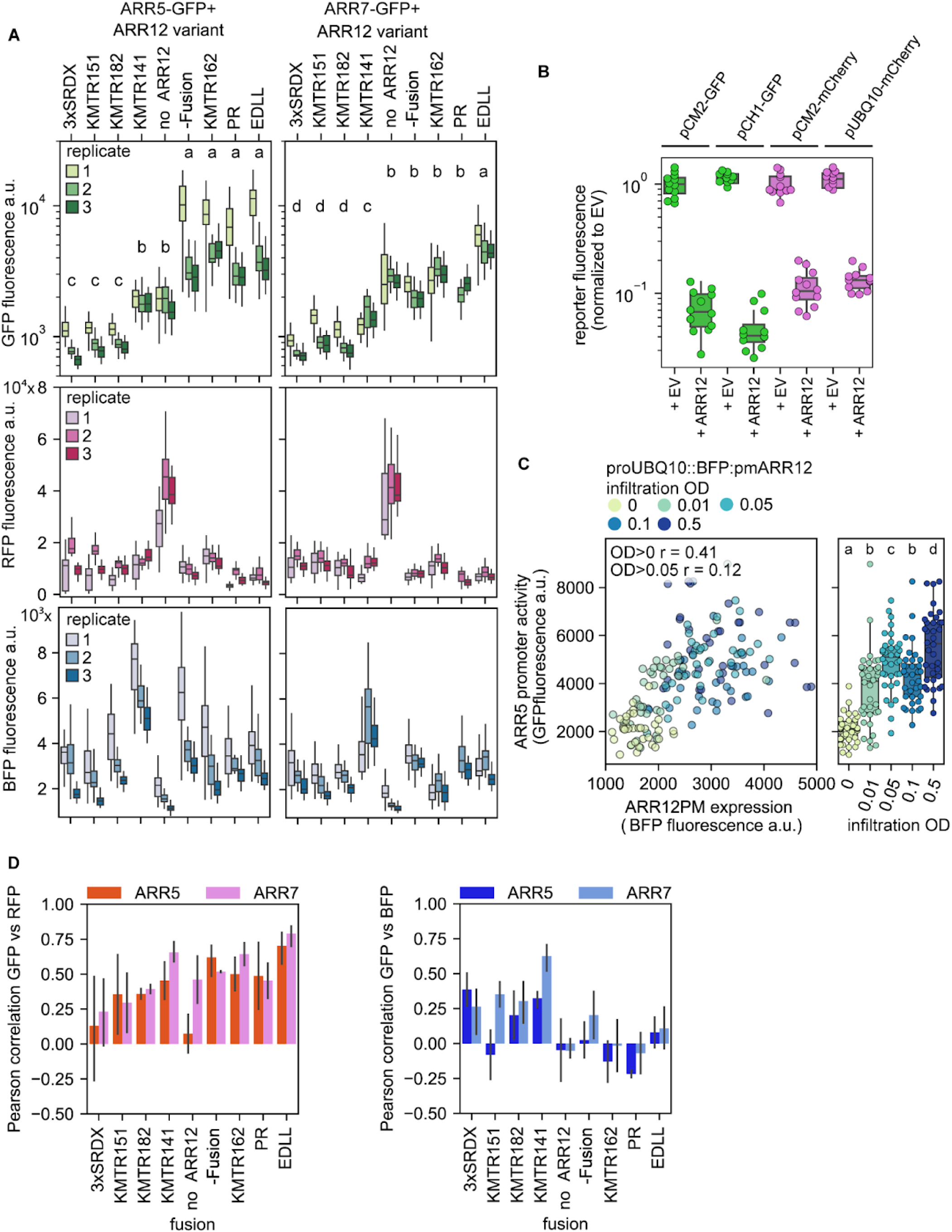
Characterization of pmARR12 variants for a range of transcriptional strength. **(A)** Top: Box plots showing the measured GFP output in *N. benthamiana* leaf discs for each BFP-pmARR12 fusion variant in 3 different replicates (n = 36 leaf disks per variant per replicate). Middle: signal in the RFP channel driven by the internal normalization strain. Bottom: signal in the BFP channel corresponding to the expression level of BFP-pmARR12 fusion variants. **(B)** Expression of ARR12 affects the normalization signal. Box plots showing the fluorescence of leaves infiltrated with GFP and RFP reporters driven by different constitutive promoters in combination with an empty vector (pCambia1300, labeled no ARR12) or the -Fusion BFP-pmARR12 variant. For each promoter, fluorescence was normalized to the mean EV fluorescence intensity. **(C)** Titration of the -Fusion BFP-pmARR12 variant shows limited concentration-dependence. The ARR12 effector strain was infiltrated at varying ODs in combination with a constant OD of the proARR5 reporter strain. Left: scatter plot of GFP vs BFP fluorescence. Right: box plots of GFP fluorescence for each ARR12 OD. **(D)** RFP, but not BFP, correlates well with reporter activity. Shown is the mean ± SD Pearson correlation between the signals of RFP and GFP (left) or BFP and GFP (right) across leaf punches for each variant and reporter combination. Box plots in (A)-(C) show the median ± IQR, whiskers show the minima and maxima excluding outliers, if present. For the GFP signal in (A), a Kruskal-Wallis test was run on each promoter dataset to determine if there were significant differences between variants, resulting in P = 1.1446e-119 for ARR5 and P = 4.1337e-112 for ARR7. The letters on top of each variant represent statistically significant groups (P < 0.05) based on Dunn’s test.

**Figure S2:**
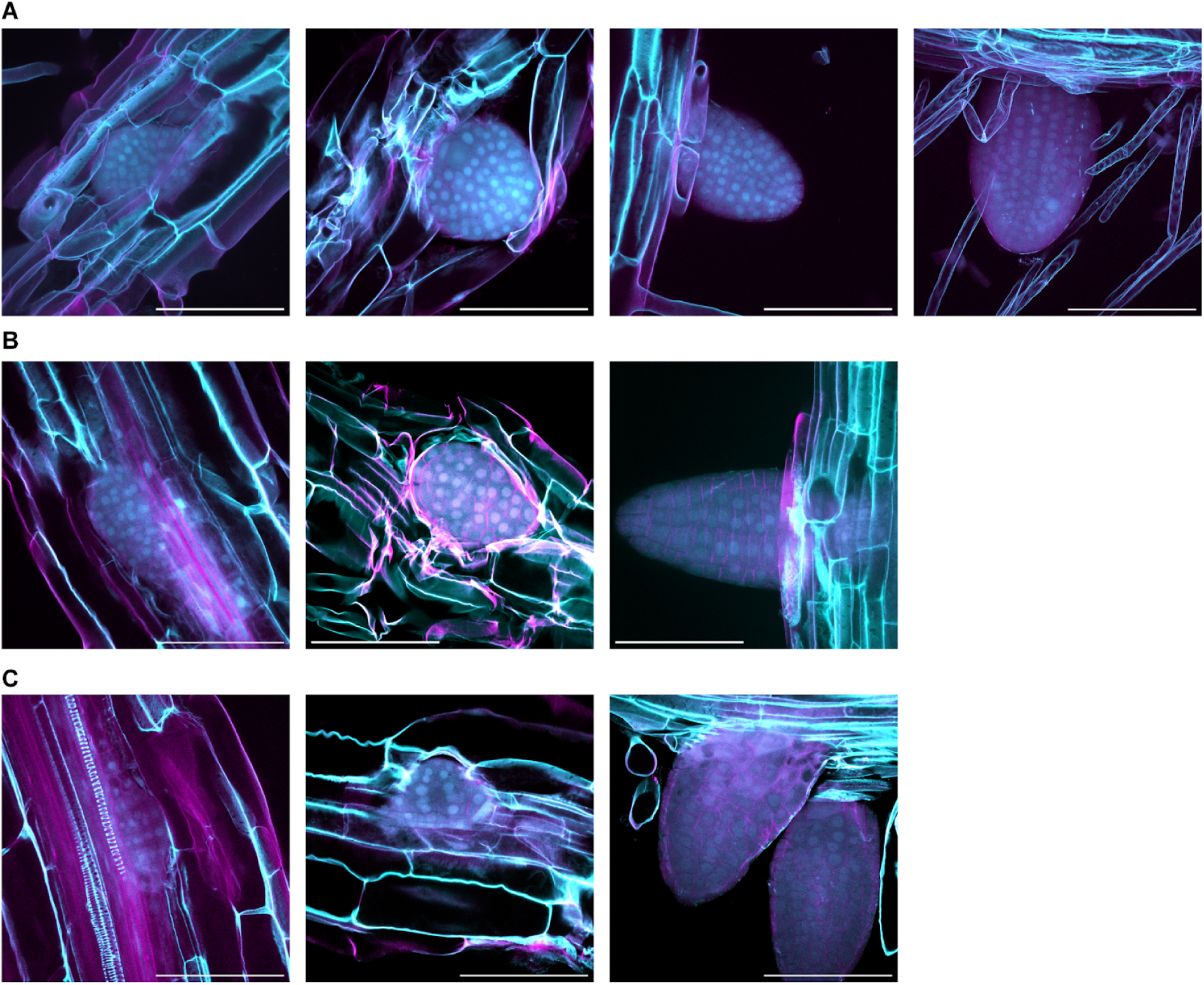
Localization of BFP-pmARR12-3xSRDX driven by the proGATA23 promoter in Arabidopsis seedlings. **(A)-(C)** Maximum projection confocal microscopy fluorescence images of proGATA23::BFP:pmARR12:3xSRDX plants. Cyan corresponds to BFP and cell wall autofluorescence. Magenta corresponds to the cell wall dye Direct Red 23. Figures (A), (B), and (C) each show details from T2 seedlings derived from T1 lines 1, 3, and 5, respectively. Scale bar = 100 µm.

